# Structural Plasticity of GABAergic Pallidothalamic Terminals in MPTP-treated Parkinsonian Monkeys: A 3D Electron Microscopic Analysis

**DOI:** 10.1101/2023.04.05.535706

**Authors:** GJ Masilamoni, H Kelly, AJ Swain, JF Pare, RM Villalba, Y Smith

## Abstract

The globus pallidus pars interna (GPi) is a major source of GABAergic inhibition upon the motor thalamus. GPi neurons are endowed with properties that allow them to fire at a high rate and maintain a tonic inhibitory influence upon thalamocortical neurons. In parkinsonism, the firing rate of GPi neurons is further increased and their firing pattern switches from a tonic to a bursty mode, two pathophysiological changes associated with increased GABAergic pallidothalamic activity. At the thalamic level, GPi terminals display ultrastructural features (large diameter, multiple synapses, large number of mitochondria) that allow them to maintain tonic synaptic inhibition at high firing rate upon thalamocortical neurons in the parvocellular ventral anterior nucleus (VApc) and the centromedian nucleus (CM), the two main GPi-recipient motor thalamic nuclei in nonhuman primates. To determine if changes of GPi neurons activity are associated with neuroplastic reorganization of GPi terminals and their synapses, we used a Single Block Facing/Scanning Electron Microscopy (SBF/SEM), high resolution 3D electron microscopic approach to compare the morphometry of GPi terminals between 2 control and 2 MPTP-treated parkinsonian monkeys. Our findings demonstrate that pallidothalamic terminals in VApc and CM undergo major ultrastructural changes in parkinsonian monkeys: (1) increased terminal volume in both nuclei, (2) increased surface area of synapses in both nuclei, (3) increased number of synapses/GPi terminals in the CM, but not VApc, (4) increased total volume of mitochondria/terminals in both nuclei but not in the number of mitochondria. In contrast, the ultrastructure of putative GABAergic terminals from the reticular thalamic nucleus was not affected in both the VApc and CM of parkinsonian monkeys. Our findings also show striking morphological differences in terminal volume, number/area of synapses and volume/number of mitochondria between GPi terminals in VApc and CM of control monkeys. In conclusion, results of this study demonstrate that GABAergic pallidothalamic terminals are endowed with a high level of structural plasticity that may contribute to the development and maintenance of the abnormal increase in pallidal GABAergic outflow to the thalamus in the parkinsonian state. Furthermore, the evidence for ultrastructural differences between GPi terminals in VApc and CM suggests that Morphologically distinct pallidothalamic terminals underlie specific physiological properties of pallidal inputs to VApc and CM in normal and diseased states.

## Introduction

The motor signs of Parkinson’s disease (PD) are mainly related to the progressive degeneration of the nigrostriatal dopaminergic (DA) pathway. The loss of striatal DA elicits changes in firing rate and pattern of neurons in the internal globus pallidus (GPi) (Filion, 1979; Miller and DeLong, 1988; Hutchison et al., 1994; Wichmann et al., 1999; Galvan et al., 2015), accompanied with an abnormal increase in GABAergic inhibition of the ventral motor thalamus, and resulting reduced thalamocortical activity (Albin et al., 1989; DeLong, 1990; Dharmadhikari et al., 2015). There is compelling evidence that the striatum and the subthalamic nucleus undergo major morphological and ultrastructural changes accompanied with robust alterations in electrophysiological and plastic properties in rodent and primate PD models of parkinsonism (Ingham et al., 1998; Day et al., 2006; Raju et al., 2008; Mathai and Smith, 2011; Villalba and Smith, 2011; Fan et al., 2012; Mathai et al., 2015; Villalba et al., 2015; Chu et al., 2017). However, despite evidence for functional and neurochemical changes of basal ganglia outputs to the thalamus in parkinsonism (see above), our understanding of the neuroplastic changes in synaptic microcircuits that could mediate these effects remains unknown. In the present study, we hypothesized that GPi terminals undergo changes in their morphology, number of synapses and mitochondrial content that may contribute to the maintenance of increased pallidal inhibition upon the ventral motor thalamus and centromedian nucleus in MPTP-treated parkinsonian monkeys.

Various sets of observations lay the foundation for this hypothesis: (1) GPi terminals display ultrastructural features (large volume, large number of mitochondria and multisynaptic innervation) tailored to maintain synaptic inhibition even at high presynaptic firing rates (Bodor et al. 2008, Wanaverbecq et al., 2008), (2) Recent findings from the subthalamic nucleus showed that increased pallidal GABAergic inhibition upon STN neurons is associated with an increased number of synapses formed by individual GABAergic terminals from the external globus pallidus (GPe) in the 6-OHDA-treated mouse model of parkinsonism (Fan et al., 2012; Chu et al., 2017), (3) The GABA release from GPi-like terminals in the thalamus and other brain regions displays low incidence of synaptic depression even when stimulated at abnormally high firing rate (Telgkamp et al., 2004; Wanavebecq et al., 2008; Rudolph et al., 2015), (4) The amount of neurotransmitter release and the number of synapses formed by multisynaptic GABAergic boutons are tightly correlated (Rudolph et al., 2015) and (5) Mitochondria are essential regulators of synaptic transmission and firing rate homeostasis since they are a major source of energy (ATP and NAD+) required for maintenance and restoration of ion gradients (Duchen, 2000; Nicholls and Budd, 2000; Toescu, 2000; Ruggiero et al., 2021). Given these observations, a detailed ultrastructural analysis of morphological changes GPi terminals undergo in the state of parkinsonism will provide a solid substrate for future studies of structure-function relationships of the pallidothalamic system in normal and diseased states.

To address this issue, we used the 3D serial block face scanning electron microscopy (SBF-SEM) approach (Ventura and Harris, 1999; Denk and Horstmann, 2004) to trace, reconstruct, and quantitatively analyze the morphometry of GABAergic pallidal terminals, synapses, and mitochondrial morphology in the basal ganglia-receiving regions of the motor thalamus in control and parkinsonian monkeys.

Results of these studies have been presented in abstract forms (Masilamoni et al., 2021a).

## Materials and Methods

### Animals

Four adult male rhesus monkeys (Macaca mulatta, 4.5–8.5 kg) from the Emory National Primate Research Center colony were used in this study (supplementary table 1). All procedures were approved by Emory’s Animal Care and Use Committee in accordance with guidelines from the National Institutes of Health. The animals were housed in a temperature-controlled room and exposed to a 12-h light/dark cycle. They were fed twice daily with monkey chow supplemented with fruits or vegetables. The animals had free access to water.

### MPTP administration and evaluation of parkinsonism

The four rhesus monkeys used in this study were divided into two groups: 2 monkeys were drug-naïve, healthy, and served as experimental controls. The other two monkeys were rendered moderately parkinsonian via chronic treatment with 1-methyl-4-phenyl-1,2,3,6-tetrahydropyridine (MPTP). The two monkeys in the MPTP treatment group received repeated injections (i.m.) of low doses of MPTP (0.2–0.7 mg/kg; Sigma-Aldrich, St Louis, USA) delivered at least one week apart until a moderate and stable state of parkinsonism emerged. The cumulative drug doses and total duration of MPTP treatment for each subject are described in supplementary table 1. The methods for evaluation of parkinsonism were described in our previous studies (Masilamoni et al., 2010; Masilamoni et al., 2011; Mathai et al., 2015; Masilamoni et al., 2021b). In brief, animals were habituated to a behavior cage equipped with infrared beams, and their spontaneous movements within this cage were monitored for 15 min weekly during the MPTP treatment period. Movements within the behavior cage were scored by an expert observer according to a nine-criteria parkinsonism rating scale, with evaluations of gross motor activity, balance, posture, arm bradykinesia, arm hypokinesia, leg bradykinesia, arm hypokinesia, arm tremor and leg tremor. Each criterion received a score of 0–3 (normal/absent to severe), for a maximal score of 30. Additionally, infrared beam breaks were counted and compared to baseline numbers measured during the pre-MPTP phase in the same subject. Animals were considered stably parkinsonian once they had achieved a score of 10 or higher on the rating scale and a >60% reduction in beam breaks from baseline, both persisting over 6 weeks following cessation of MPTP treatment. The final rating scores for the two MPTP-treated subjects in this study ranged from 10 to 14, corresponding to moderate parkinsonism.

### Anterograde labeling of pallidothalamic terminals

In the two control and two MPTP-treated monkeys, pallidothalamic terminals were labeled anterogradely with viral vector injections in the GPi (Swain et al., 2020). A total of 2–8 μL of AAV5-hSyn-ChR2-EYFP or AAV5-hSyn-Arch3-EYFP was delivered in the GPi. In the control monkey, the injection was made with a microsyringe into the GPi, while the animal’s head was fixed in a stereotaxic frame. The surgical procedure was achieved under isoflurane anesthesia and sterile conditions. Following the procedure, the animals received daily injections of analgesics for 72 hours to reduce pain. Pre-operative MRI scans helped delineate the stereotaxic coordinates. In the MPTP-treated monkeys, recording chambers were stereotactically directed at the pallidum on both sides of the brain and attached to the skull. After surgery, electrophysiological mapping was conducted to help define the borders of the GPe and GPi. GPi cells were identified based on the depth of the electrode (at least 2 mm ventral to the first GPe unit), the presence of “border” cells between GPe and GPi (DeLong, 1971), and the presence of neurons that fired at high frequency, characteristic for GPi cells (DeLong, 1971; Galvan et al., 2005; Galvan et al., 2011). The injection was delivered into the GPi using a probe combined with a recording microelectrode (Kliem and Wichmann, 2004). Extracellular recordings were performed while lowering the microsyringe to determine the final location in the GPi for the injection (Galvan et al., 2010; Kliem et al., 2010). Both monkeys survived at least 6 weeks after the virus injections to allow efficient anterograde transport to GPi terminals in the thalamus.

### Tissue collection and processing for microscopy

At the completion of the study, the animals were euthanized with an overdose of pentobarbital, and transcardially perfused with a Ringer’s solution and a mixture of paraformaldehyde (4%) and glutaraldehyde (0.1%). The brains were removed from the skull, postfixed in 4% paraformaldehyde, and cut in serial sections (60 µm) with a vibratome. The sections were stored at −20°C until further histological processing. Thalamic tissue sections containing the basal ganglia (BG)-receiving parvocellular ventral anterior (VApc) or the centromedian (CM) nuclei, as delineated in the rhesus monkey stereotaxic brain atlas (Paxinos et al. 1999) and in adjacent calbindin-immunostained sections (Calzavara et al. 2005), were removed from the anti-freeze solution and placed in phosphate-buffered saline (PBS, 0.01 M, pH 7.4). Sections were prepared for electron microscopy and placed in a cryoprotectant solution (phosphate buffer [PB]= 0.01 M at pH 7.4, with 25% sucrose and 10% glycerol) for 20 minutes, frozen at -80°C for 20 minutes, thawed, and returned to a graded series of cryoprotectant solution (100%, 70%, 50%, 30%) diluted in PBS. Sections were washed in PBS and then pre-incubated in a solution of 10% normal goat serum and 1% bovine serum albumin in PBS for 1 hour. Sections were immunostained for Green Fluorescent Protein (GFP) to localize anterogradely labeled GPi terminals in the VApc and CM. This tissue was incubated with a primary rabbit antibody (1:5,000 dilution) for 48 hours at 4°C. Next, sections were rinsed in PBS and transferred for 1.5 hours to a solution with a secondary biotinylated goat anti-rabbit antibody (1:200 dilution). After, sections were placed in a solution of 1% avidin-biotin-peroxidase complex (Vector Laboratories, Burlingame, CA USA), washed in PBS and Tris buffer (0.05 M, pH 7.6), and transferred to a solution containing 0.01M imidazole, 0.005% hydrogen peroxide, and 0.025% 3,3′-diaminobenzidine tetrahydrochloride (DAB; Sigma, St. Louis, MO) in Tris for 10 minutes. Several rinses of the tissue in PBS ended the DAB reaction. Sections with the maximum amount of GFP immunostaining in VApc and CM were put into vials in phosphate buffer solution and sent to Cleveland Clinic (Cleveland, OH) in 4% paraformaldehyde for SBF-SEM processing. In preparation for the SBF-SEM, the tissue went through a multi-day staining process beginning with washing off aldehydes in sodium cacodylate buffer (0.1M). The tissue samples were then incubated in 1.5% potassium ferrocyanide and 2% osmium tetroxide (in sodium cacodylate buffer, 0.1M) at 4°C, rinsed in double distilled H_2_O (dd H_2_O), and incubated in 1% thiocarbohydrazide at 60°C, followed by washes in ddH_2_O. Next, the tissue was placed in 2% osmium tetroxide on a rotator at RT, rinsed in ddH_2_O, and left at 4°C for 48-72 hours in a saturated aqueous solution of uranyl acetate and then lead stained at 60°C, rinsed in ddH_2_O, and dehydrated in a series of graded ethanol and propylene oxide. Finally, the tissue was placed in the resin (EMbed-812), first a mixture of resin and propylene oxide (50/50) and after that, in fresh resin (100%) and embedded in Pelco silicone mold at 60°C until the resin was fully cured (8-14 hours). Next, the resin blocks were removed from the mold, trimmed, and mounted on an aluminum pin where the vertical sides of the sample were covered with colloidal silver liquid. Once the imaging surface of the tissue was exposed, the sample was then set up in the SBF-SEM in-chamber ultramicrotome and examined in the electron microscope. Low, medium, and high-resolution 2D images were initially taken of the block face surface to assess the tissue preservation. Multiple series of 200-300 EM images were then obtained using two SBF-SEM systems: a Zeiss Sigma VP scanning EM with Gatan 3View system, and a ThermoFisher Teneo VolumeScope.

## 3D Reconstruction from Serial Sectioning Electron Microscopy

Selected areas in the VApc and CM with dense GFP-labeled GPi terminals were used for 3D reconstruction employing the SBF/SEM method. Approximately 200-400 serially scanned micrographic images (∼70 nm-thick) were collected from each region of interest and labeled and unlabeled GPi terminals were chosen at random from these images to be reconstructed using the 3D software Reconstruct (available at: synapses.clm.utexas.edu). In order to avoid bias in the selection of elements being reconstructed, two experimenters, one of them blinded to the treatment (control vs MPTP treated) reconstructed and analyzed individual terminals. Identification of axon terminal subtypes in the VApc and CM was based on ultrastructural features previously reported in EM studies of the mammalian motor thalamus (Ilinsky et al., 1997; Kultas-Ilinsky et al., 1997; Jones, 2002) and in an EM study from our laboratory (Sidibe et al., 1997; Swain et al., 2020). Small (i.e. ∼0.5-0.7 μm in diameter) terminals forming asymmetric synapses were categorized as originating from the cerebral cortex; medium-to-large sized (∼0.5-1.5 μm in diameter) terminals forming single symmetric synapses were considered as GABAergic terminals from the thalamic reticular nucleus (RTN type); and large terminals enriched in mitochondria (∼1-3 μm in diameter) forming multiple symmetric synapses with single postsynaptic targets were categorized as putative GABAergic terminals from the GPi. To further confirm that GPi was the source of terminals forming multiple symmetric synapses, AAV5-eYFP was injected into the GPi of 2 control and 2 MPTP-treated monkeys (see above), and the morphology of 40 anterogradely labeled terminals was examined in the present study. As expected, anterogradely labeled GPi terminals were large (1.0–3.0 μm in diameter), densely filled with mitochondria, and formed multiple axodendritic synapses onto dendrites in VApc (Figure 1A, B) and CM (Figure 1C, D). Using these structural criteria, we considered 40 unlabeled large multisynaptic terminals in the VApc and CM as putative GPi terminals and added those to the sample of GPi boutons reconstructed in this study. TIFF images of single sections were imported into Reconstruct and calibrated with the section thickness and pixel size provided by Renovo. Finally, labeled and unlabeled terminals, mitochondria, synapses, and dendrites from each object analyzed were manually traced in each serial electron micrograph using Reconstruct (Villalba and Smith, 2010, 2011). From these serially identified elements, the software created a 3D representation of each object from which it calculated the volume of terminals, mitochondria, and the surface area of synapses. The number of mitochondria and synapses were also recorded in control and MPTP-treated monkeys. Only terminals that could be seen through their full extent in serial sections were reconstructed; a series of 30-100 scanned ultrathin images were used depending on the size of terminals. From GFP-immunostained VApc sections, 10 randomly chosen labeled and unlabeled GPi terminals from 2 normal and 2 MPTP-treated monkeys were used for 3D reconstruction. Further, we used the mitochondrial complexity index (MCI) to quantify changes in mitochondrial shape complexity (Vincent et al., 2019; Faitg et al., 2021).

**Figure. 1.**
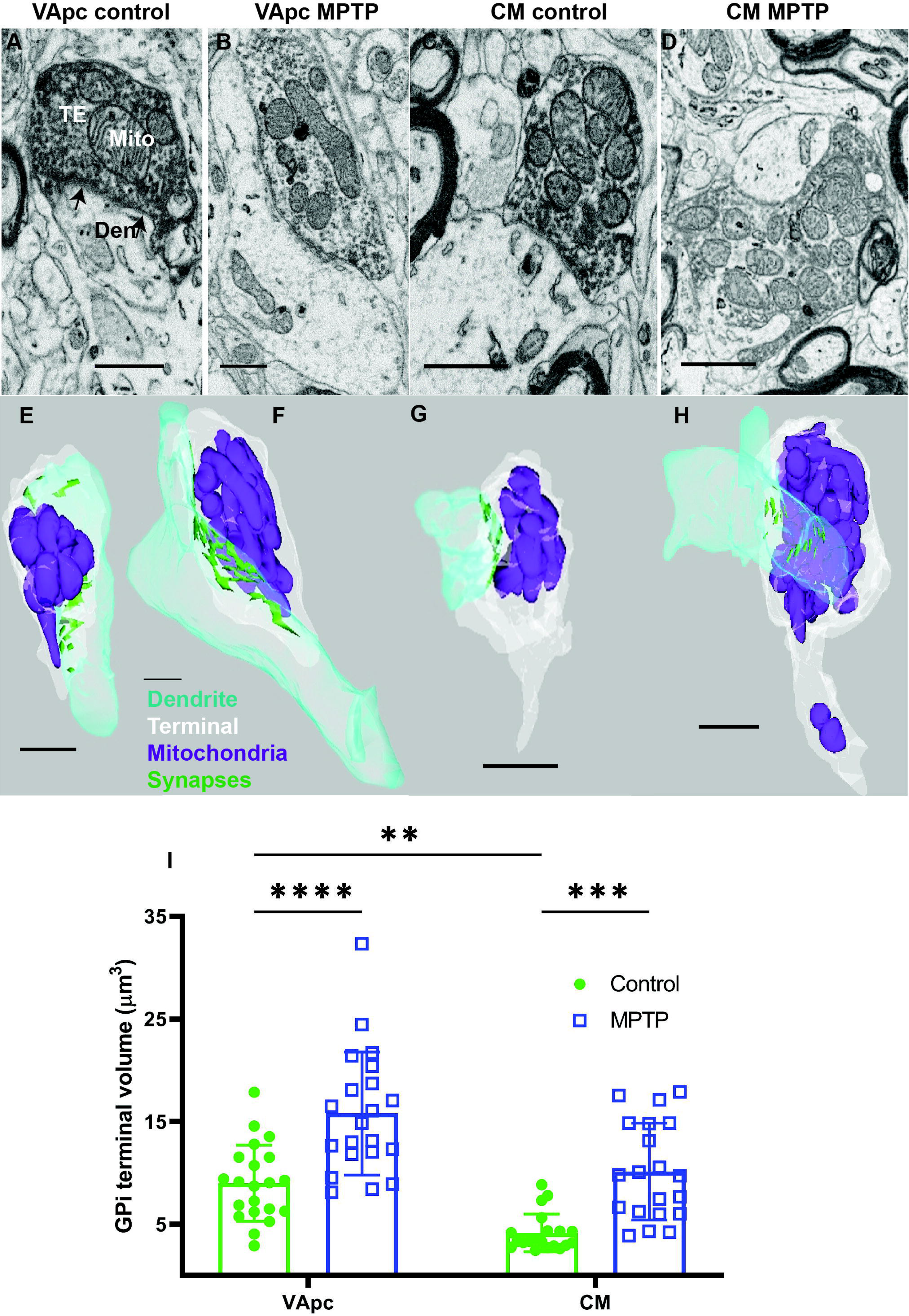
Electron micrographs (A-D) and corresponding 3D EM reconstructions (E-H) of GFP-positive (A,C) or negative (B,D) GPi terminals (Te) that form multiple symmetric axo-dendritic synapses in the VApc (A,B) and CM (C,D) of control or MPTP-treated parkinsonian monkeys. Abbreviations: Den (dendrite), M (mitochondria). Scale bars: A–H = 1μm. (I) Histogram comparing the relative volume of GPi terminal in the VApc and CM between control and MPTP-treated parkinsonian monkeys. Each data point is a terminal (n = 20 per group). Statistical differences were determined by two-way ANOVA for repeated measures followed by the Tukey post hoc test. Significance was taken at PLJ<LJ0.05*, PLJ<LJ0.001**, and PLJ<LJ0.0001***. All results are expressed as meanLJ±LJstandard deviation (SD).

### Statistical Analysis

Data were statistically analyzed using GraphPad Prism software (version 9.3). Multiple comparisons for two-way ANOVA for repeated measures followed by the Tukey post hoc test was used to compare terminal volume, surface area of synapses, mitochondria volume, and number of synapses or mitochondria per terminal, between control and MPTP treatments. Significance was taken at PL<L0.05*, PL<L0.001**, and PL<L0.0001***. All results are expressed as meanL±Lstandard deviation (SD).

## Results

### Nigrostriatal dopamine denervation in MPTP-treated monkeys

Chronic low dose MPTP exposure was used in the present study to slowly induce progressive parkinsonian motor signs and nigrostriatal dopaminergic denervation in two monkeys. A detailed description of this MPTP treatment protocol, quantitative data about parkinsonian motor scores, extent of striatal dopamine denervation, and nigral dopaminergic neurons loss are provided in our previous study (Swain et al., 2020). In brief, the post-commissural and lateral edge of the pre-commissural putamen exhibited the most severe reduction in tyrosine hydroxylase (TH) immunoreactivity, followed by the head and body of the caudate nucleus, which were also significantly affected, while immunoreactivity was much less reduced in the nucleus accumbens (Figure S1). At the midbrain level, the ventral tier of the substantia nigra pars compacta (SNc) was severely damaged, whereas a significant number of TH-immunoreactive neurons and processes remained in the ventral tegmental area and dorsal tier SNc (Swain et al., 2020). These results are consistent with previous findings from our laboratory using the same animal model (Masilamoni et al., 2010; Masilamoni et al., 2011).

### Pallidothalamic terminals in VApc and CM undergo robust ultrastructural changes in MPTP-treated parkinsonian monkeys

Pallidothalamic terminals in VApc and CM of control and MPTP-treated monkeys were first identified as GFP-labeled terminals forming synaptic contact with projection neurons (no vesicle in the postsynaptic targets). As expected, anterogradely labeled GPi terminals were large (1.0–3.0 μm in diameter), enriched in mitochondria, and involved in the formation of multiple axodendritic synapses onto dendrites in VApc (Figure 1A, B) and CM (Figure 1C, D). We randomly selected and reconstructed 80 individual GPi terminals out of 2 control (VApc: 20, CM: 20) and 2 parkinsonian (VApc: 20, CM: 20) animals, respectively with an x,y resolution of 7 nm and a z resolution of 50 nm (30-100 serial images per GPi terminal). Each pallidothalamic terminal was manually traced in its full extent through a series of images which were then uploaded in the Reconstruct software to build 3D structures (Figure 1E–H). From these reconstructed terminals, we measured and compared their volume between VApc and CM in control and parkinsonian monkeys. This analysis revealed a significant increase in the average volume of individual GPi terminals in the VApc and CM of MPTP-treated monkeys relative to controls (Figure 1I; p <.0001, t test). Furthermore, our 3D quantitative analysis demonstrated that the GPi terminal volume in the control VApc (mean 8.99µm^3^) was significantly larger than in CM (P<0.0001; mean 4.14 µm^3^).

### Morphometric changes of pallidothalamic synapses in parkinsonian monkeys

The number, size, and shape of synapses are key structural determinants of synaptic efficacy, strength, short-term, and long-term plasticity (Matz et al., 2010; Sudhof, 2012; Wilhelm et al., 2014; Holler et al., 2021). To determine if the morphometry of pallidothalamic synapses was altered in parkinsonian monkeys, we used the 3D SBF-SEM reconstruction approach to quantify the number and surface area of symmetric synapses formed by individual GPi terminals in controls and MPTP-treated parkinsonian monkeys. From the 80 reconstructed pallidal terminals described above (Figure 1), a total of 399 (266/133 in control VApc/CM) and 595 (277/318 in parkinsonian VApc/CM) synapses were morphometrically analyzed. The total number and surface area (SA) of synapses formed by individual GPi terminals were significantly increased in CM of parkinsonian monkeys (Figure 2A, B) (P<0.0001 and P<0.0001 respectively). However, there was no significant difference in the number and SA of synapses in the VApc between normal and MPTP-treated monkeys (Figure 2A, B). In controls, the median number of synapses established by single terminals was 13.3 in VApc (n=20; minimum to maximum, 5–35) and 6.65 in CM (n=20; minimum to maximum, 3–12; Figure 2A, B). In each case, all synapses from each terminal converged on a single postsynaptic target.

**Figure 2.**
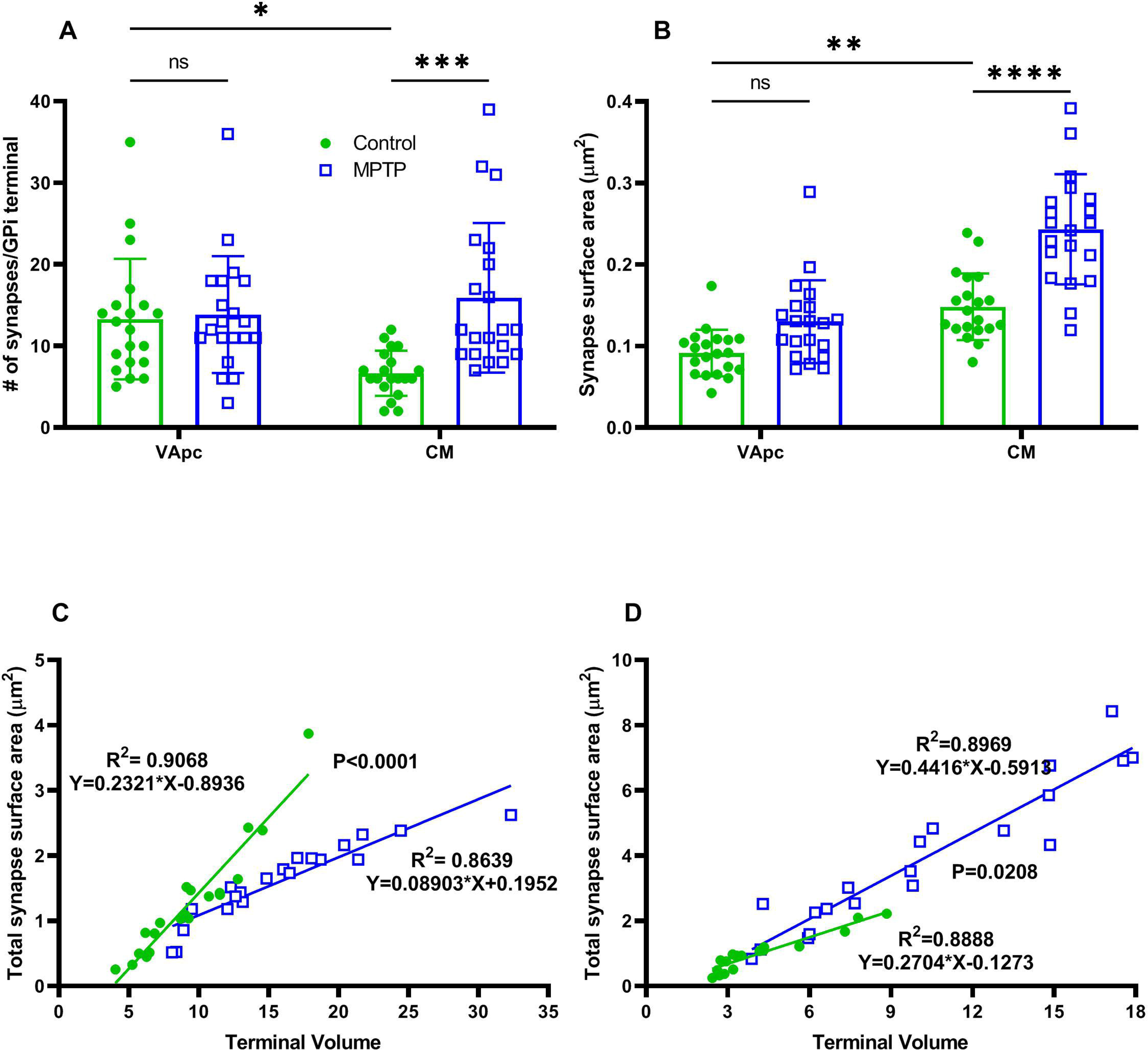
(A-B) Comparison of synapses number and surface area of GPi terminals in VApc and CM between control and parkinsonian monkeys. No significant difference was found in number of synapses and surface area of GPi terminals in VApc between control and parkinsonian animals. The number of synapses and surface area per GPi axon terminals are significantly larger in parkinsonian monkeys than in controls (one-way ANOVA with Tukey’s post hoc test, ***P = 0.0005; ****P < 0.0001). GPi terminals in VApc have a greater number of synapses than in CM (P=0.02; A), but the synapses of GPi terminals in CM have a larger surface area than in VApc (P<0.002; B). (C-D) Scatter diagrams showing the linear regression analysis of terminal volume (μm^3^) and total surface area (μm^2^) for control (green; n=20) and parkinsonian monkeys (blue; n=20) in VApc (C) and CM (D). In all cases, the terminal volume was positively correlated with the total synapses surface area. Significant differences in slopes were found between the control and parkinsonian monkeys in both VApc and CM (P<0.0001, P=0.0208; C-D).

Our 3D EM analysis also revealed a significant difference in the number and SA of synapses between the VApc and CM of control monkeys such that GPi terminals in the VApc harbored a larger number of synapses than in CM (P<0.05), but pallidothalamic synapses in CM have a larger SA than in VApc (p<0.001) (Figure 2A, B). These structural differences were not found in parkinsonian monkeys. To determine if the morphology and prevalence of synapses were related to the size of GPi terminals, correlations were made between the total number or SA of synapses and the volume of GPi terminals. Positive correlations were found between the GPi terminal volume and the SA of synapses in both VApc and CM of control and parkinsonian monkeys (Figure 2C, D). However, the regression lines for control and MPTP-treated group have significantly different intercepts and slopes in VApc (Figure 2C, P<0.0001), but not so much in CM (Figure 2D, P=0.0208).

### Mitochondrial morphology is significantly altered in pallidothalamic terminals of parkinsonian monkeys

Mitochondria in axon terminals are critical for the mobilization of the reserve pool of synaptic vesicles and for the regulation of synaptic strength (Shepherd and Harris, 1998; Rowland et al., 2000; Billups and Forsythe, 2002; Verstreken et al., 2005; Gazit et al., 2016; Smith et al., 2016; Cserep et al., 2018). Alterations in mitochondrial size, shape and number are frequently encountered in neurological diseases (Whiting et al., 1979; Mortiboys et al., 2008; Guo et al., 2017; Trinh et al., 2021). GPi terminals are enriched in mitochondria (Figure 1). Because of the high and tonic firing rate of GPi neurons, the neurotransmitter release and neuroplastic properties of pallidothalamic terminals are highly dependent on a constant and reliable supply of energy through mitochondrial respiration. Given recent evidence that changes in mitochondrial morphology and prevalence may contribute to the pathophysiology of brain disorders (Trimmer et al., 2000; Brustovetsky et al., 2021; Liu et al., 2021; Toomey et al., 2022), we used the 3D SBF/SEM approach to compare mitochondrial morphology in pallidothalamic terminals in VApc and CM between control and parkinsonian monkeys.

All mitochondria from the 80 (40 control and 40 parkinsonian) GPi terminals analyzed in this study were fully reconstructed and morphometrically characterized. A total of 285 (173/122 in control VApc/CM) and 398 (220/178 in parkinsonian VApc/CM) mitochondria were manually traced from each image stack to generate 3D reconstructions. Data revealed that the mitochondrial volume was significantly increased in the VApc (p = 0.0001) and CM (p = 0.0001) of parkinsonian monkeys compared with controls (Figure 3A). On average, pallidothalamic terminal mitochondria in VApc and CM were 175% and 259% larger in parkinsonian monkeys, respectively (mean volume = 3.98 µm^3^ and 2.55 µm^3^), than in controls (mean volume = 2.26 µm^3^ and 0.98 µm^3^). However, no significant difference was found in the number of mitochondria/terminals between control and MPTP-treated monkeys (Figure 3B). Mitochondrial volume heterogeneity was significant in VApc and CM (Figures 3A, C, D). Frequency distribution analysis showed that ∼35% of mitochondria in GPi terminals of parkinsonian monkeys were large (volume of 0.6 – 1.3 µm^3^ and 0.4-1.0 µm^3^ VApc and CM, respectively, Figures 3C, D), while only 8% in VApc and 4% in CM were within these volume ranges in control monkeys. To determine whether the volume of mitochondria was related to overall size of GPi terminals, correlation analyses were performed. Strong positive correlations were found between GPi terminal volumes and mitochondrial volumes (R^2^=0.7258 and R^2^=0.4656 VApc and CM, respectively; Figure 3E, F) in both control and parkinsonian monkeys. Notably, the regression lines for control and MPTP-treated group have no significant difference between the intercepts and slopes (P=0.2718, P=0.8109, VApc and CM, respectively) (Figure 3E, F). Further, our data demonstrated a positive correlation between the mitochondrial volume and the SA of synapses in both VApc and CM of controls and parkinsonian monkeys (Figure 3G, H). The regression lines for control and MPTP-treated group have significantly different intercepts and slopes in VApc (Figure 3G, P<0.0012), but not in CM (Figure 3H, P=0.1237).

**Figure 3.**
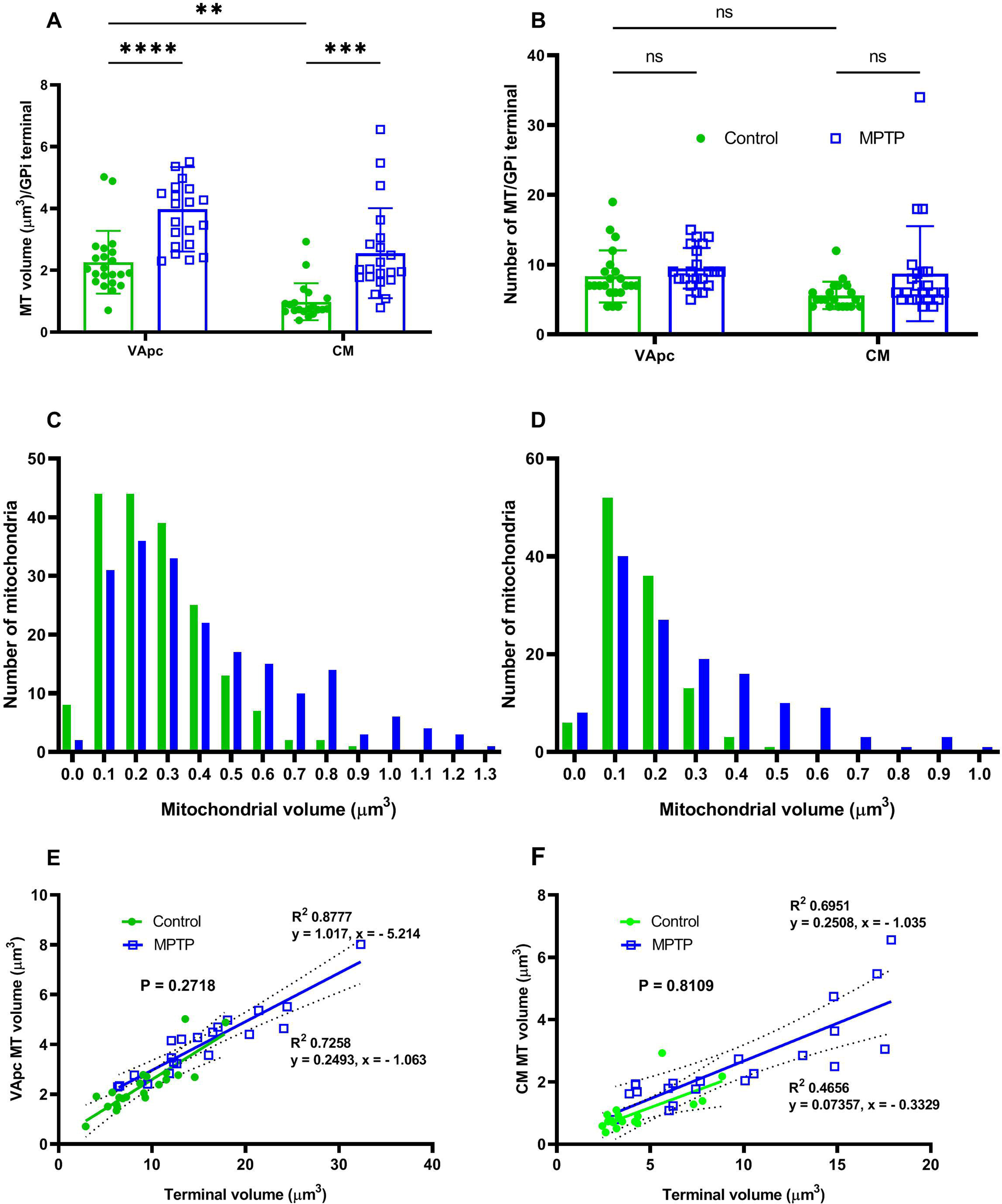
(A-B) Histograms comparing morphometric measurements (volume, and number) of mitochondria in 3D-reconstructed GPi terminals of VApc and CM between control and parkinsonian monkeys. The mitochondrial volume per GPi terminals in the VApc and CM was significantly larger in MPTP-treated monkeys compared with controls (one-way ANOVA with Tukey’s post hoc test; ****P < 0.0001; ***P = 0.0003; A). The mitochondrial volume/GPi terminals was larger in VApc than in CM (**P = 0.0039). No significant difference was found in the number of mitochondria/GPi terminals in the VApc and CM between control and parkinsonian monkeys (B). (C-D) Histograms comparing the relative frequency distribution of mitochondrial volume in GPi terminals of VApc (C) and CM (D) between controls and parkinsonian monkeys. Total number of mitochondria used for control [n = 275 (VApc: 173 and CM = 112)] and MPTP-treated monkeys. [n = 398 (VApc: 220 and CM :178)]. (C-D) Histograms comparing the relative distribution of mitochondrial volumes in GPi terminals of VApc and CM between control and parkinsonian monkeys. Note the higher proportion of large mitochondria in both nuclei of parkinsonian animals. (E-F) Scatter diagrams showing the linear regression analysis of terminal volume (μm^3^) and total mitochondria volume (μm^3^) for control (green; n=20) and MPTP-treated (blue; n=20) monkeys in VApc (E) and CM (F). In all cases, the terminal volume correlated positively with the total mitochondria volume (E and F). (G-H) Scatter diagrams showing the linear regression analysis of mitochondria volume (μm^3^) and total surface area (μm^2^) for control (green; n=20) and MPTP-treated (blue; n=20) monkeys in VApc (G) and CM (H). In all cases, the mitochondrial volume correlated positively with the total surface area (G and H).

### Mitochondrial complexity index and mitochondrial volume density between control and parkinsonian monkeys

To further assess potential changes in mitochondrial morphology of pallidothalamic terminals between control and parkinsonian condition, we measured the mitochondrial complexity index (MCI) (Vincent et al., 2019; Faitg et al., 2021). Based on this metric, mitochondria in GPi terminals of the VApc of parkinsonian monkeys were significantly more complex than those of controls (mean MCI = 1.351 and 0.8924, unpaired t test P<0.0065, Figures 4A). However, such was not the case in CM (P = 0.139, Figure 4A). No MCI difference was found between VApc and CM of control and parkinsonian animals (Figure 4A). Next, we determined the relationships between the two variables of interest, i.e., individual mitochondrial volume and its corresponding MCI, together reflecting the mitochondrial phenotype or mitotype of GPi terminals in VApc (Figure 4C) and CM (Figure 4D) of control and parkinsonian monkeys. Results of this regression analysis demonstrated significant dissimilarities in the mitochondrial morphology between control and MPTP-treated monkeys, in both VApc and CM (P=0.0001 and P=0.0001, Figure 4C, D). They also showed that the regression lines for control and MPTP-treated monkeys have significantly different intercepts and slopes (Figure 4C, D).

**Figure 4.**
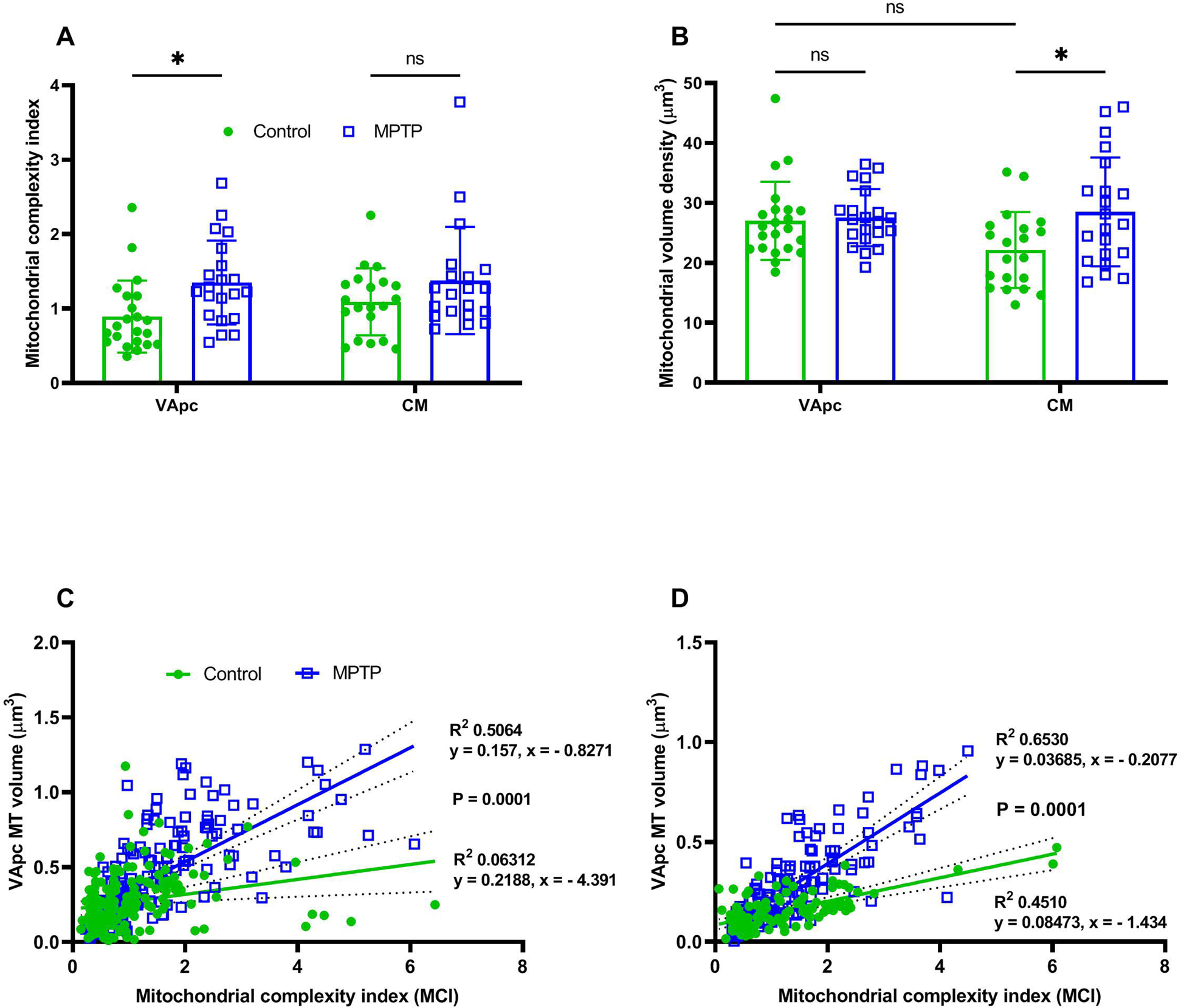
(A-B) Histograms comparing the morphological measurements of Mitochondrial complexity index (MCI; calculated using the equation: SA^3^/16π^2^V^2^, where SA is average mitochondrial surface area within a terminal and V is the average mitochondrial volume within the terminal) (A) and the mitochondrial volume density (MVD; mitochondrial volume normalized to terminal volume) (B) of mitochondria in 3D-reconstructed GPi terminals of VApc and CM between control and parkinsonian monkeys. The MCI and MVD are significantly larger in VApc and CM respectively, of parkinsonian monkeys than in controls (one-way ANOVA with Tukey’s post hoc test, *P < 0.02; *P < 0.01). No significant difference was found in MCI and MVD in CM and VApc respectively, between control and parkinsonian animals. (C-D) Plotting individual mitochondrial volume and their corresponding MCI together on a mitochondrial phenotype (i.e., mitotype) graph for control (green; n=186 and 112) and parkinsonian monkeys (blue; n=197 and 136) in VApc (C) and CM (D) respectively, highlights significant mitochondrial morphological dissimilarities between control and parkinsonian monkeys (P <0.0001).

To understand how MPTP-induced changes in terminal volume relate to changes in mitochondrial morphology, we measured mitochondrial volume density (MVD). MVD is defined as the percentage of terminal volume occupied by mitochondria, in 3D reconstructed models (Vincent et al., 2019; Faitg et al., 2021). The data suggest that there was a significant increase (30.3%; P = 0.0234) in the MVD within GPi terminals in CM, but not in VApc, of parkinsonian monkeys (Figure 4B). No significant change in MVD was found between the VApc and CM of control and parkinsonian groups (Figure 4B).

### Morphometric analysis and mitochondrial content of RTN-like GABAergic terminals in parkinsonian monkeys

To determine whether the morphological and mitochondrial changes found in GPi terminals were also seen in other GABAergic terminals in VApc and CM, we reconstructed and analyzed the morphology and mitochondrial content of 90 (22/21 in control VApc/CM and 26/21 parkinsonian VApc/CM) RTN-like GABAergic terminals in the VApc and CM of parkinsonian and control monkeys (Figure 5A–H). From these terminals, a total of 135 mitochondria (27/40 in control VApc/CM and 34/34 in parkinsonian VApc/CM) were manually traced from each image stack and reconstructed. As expected, 3D-reconstructed RTN-like terminals were much smaller in size and displayed a lower mitochondrial volume than GPi terminals in VApc and CM (compare Figures 1I, J and 3A with figure 5I, J). In contrast to GPi terminals, no significant difference in terminal volume and mitochondrial volume was found in RTN-like terminals of VApc and CM between the control and parkinsonian states (Figure 5I, J).

**Figure. 5.** Electron micrographs (A-D) and reconstructed 3D models (E-H) of RTN-like terminals (Te) that form single symmetric axo-dendritic synapses in the VApc (A, B) and CM (C, D) of control and MPTP-treated parkinsonian monkeys. Abbreviations: Den (dendrite), M (mitochondria). Scale bars: A=1 um (valid for B-D); E= 1μm (valid fir F-H) I: Histogram comparing the relative volume of RTN-like terminals in the VApc and CM of controls and parkinsonian monkeys. Each data point is a terminal (n = 20 per group). Statistical differences were determined by two-way ANOVA for repeated measures followed by the Tukey post hoc test. No ultrastructural differences were noticed in putative RTN GABAergic terminals in VApc and CM of MPTP-treated monkeys.

## Discussion

In this study, we used the SBF-SEM 3D electron microscopy approach to assess ultrastructural changes in the morphometry, synaptic connections, and mitochondrial content of GPi terminals in the basal ganglia-receiving regions of the ventral motor thalamus and the CM of MPTP-treated parkinsonian monkeys. Our findings demonstrate that pallidothalamic terminals in VApc and CM undergo major ultrastructural changes in parkinsonian monkeys: (1) increased terminal volume in both nuclei, (2) increased surface area of synapses in both nuclei, (3) increased number of synapses/GPi terminals in the CM, but not VApc, (4) increased total volume, but not number, of mitochondria/terminals in both nuclei. In contrast to GPi terminals, the ultrastructure of putative GABAergic terminals from the reticular thalamic nucleus was not affected in both the VApc and CM of parkinsonian monkeys. Despite a common cellular origin, our findings also show that GPi terminals in VApc and CM display different ultrastructural features, raising the possibility that pallidal inputs differentially regulate neuronal activity in these two thalamic nuclei. Together, these findings demonstrate that the pallidothalamic system undergoes significant structural neuroplastic changes in the diseased state, suggesting a potential contribution of disrupted structure-function relationships of the pallidothalamic system in parkinsonism.

### Anatomical and functional characteristics of pallidothalamic terminals in VApc and CM of non-human primates

In keeping with our previous study (Sidibe et al., 1997) and others (Parent and De Bellefeuille, 1983; Ilinsky and Kultas-Ilinsky, 1987; Francois et al., 1988; Fenelon et al., 1990; Hazrati and Parent, 1991; Rouiller et al., 1994; Sakai et al., 1996), injections of AAV5-GFP in the ventrolateral GPi resulted in dense anterograde labelling in the sectors of the VA/VL and CM known as the source of projections to motor cortices and the sensorimotor striatum (Ilinsky and Kultas-Ilinsky, 1987; Matelli et al., 1989; Darian-Smith et al., 1990; Zemanick et al., 1991; Inase and Tanji, 1995; McFarland and Haber, 2002; Sidibe et al., 2002; Smith et al., 2014).

Consistent with the previous literature (Grofova and Rinvik, 1974; Kultas-Ilinsky et al., 1983; Smith et al., 1987; Smith et al., 1994; Sidibe et al., 1997; Rovo et al., 2012; Swain et al., 2020), our 3D EM reconstruction data demonstrate that pallidothalamic terminals were large (1.0–3.0 μm in diameter), densely filled with mitochondria, and formed multiple axodendritic synapses in the VApc and CM of rhesus monkeys. Combined anatomical and physiological data have revealed the presence of other large, multisite GABAergic terminals from the zona incerta (Bartho et al., 2007) and the anterior pretectal nucleus (APT) (Bokor et al., 2005) that exert strong inhibitory effects upon their postsynaptic thalamic targets (Bartho et al., 2002; Bokor et al., 2005; Lavallee et al., 2005; Rovo et al., 2012; Halassa and Acsady, 2016). It has been suggested that the ultrastructural features of APT terminals enable them to exert strong inhibitory influences on neuronal activity and maintain tonic synaptic transmission at high presynaptic firing rates in the posterior nucleus of the rodent thalamus (Xu-Friedman and Regehr, 2004). Activation of the multi-synaptic APT terminals, indeed, generated a larger charge transfer and greater persistent current, even at high stimulation frequencies, in thalamocortical cells (Wanaverbecq et al., 2008). In contrast, the monosynaptic GABAergic RTN terminals failed to maintain tonic inhibition (Wanaverbecq et al., 2008), thereby suggesting that the morphology and synaptic architecture or GABAergic pallidothalamic afferents may dictate their strength and neuroplastic properties in normal and parkinsonian conditions (see details below).

The present 3D volumetric analysis confirmed that GPi terminals in the VApc and CM display characteristically different ultrastructural features in control monkeys (Sadikot et al., 1992; Balercia et al., 1996; Sidibe et al., 1997). Our findings show striking morphological differences in number/area of synapses and volume/number of mitochondria between GPi terminals in VApc and CM of normal monkeys. Given that pallidothalamic axonal projections to the VApc and CM originate from the same GPi neurons (Parent and De Bellefeuille, 1983; Parent and Hazrati, 1995; Sidibe et al., 1997; Sidibe et al., 2002), these observations suggest target-specific ultrastructural differences in the morphometry of GPi terminals between the VApc and CM in nonhuman primates. Whether such anatomical differences are reflected in the synaptic neuroplastic properties and strength of pallidothalamic synapses that impinge upon CM vs VApc neurons in normal and diseased states remain to be established. Such studies could provide important information about possible functional consequences of morphometric differences of pallidothalamic terminals and their associated synaptic microcircuitry in the basal ganglia-receiving regions of the motor and intralaminar thalamus in normal and parkinsonian states.

### Morphometric changes of pallidothalamic terminals and synapses in parkinsonian monkeys: Potential functional significance

Our findings demonstrate that the volume as well as the number and size of synapses formed by pallidothalamic terminals in VApc and CM undergo robust ultrastructural changes in MPTP-treated parkinsonian monkeys, whereas the morphology of RTN GABAergic terminals remain intact under these conditions (Figure 5A–J). Given converging evidence that large terminal volume and multisynaptic connections have an impact upon neurotransmitter release, synaptic strength and synaptic plasticity in the mammalian thalamus (Bodor et al., 2008; Nishijima et al., 2020), it is tempting to speculate that the increased volume of GPi terminals in parkinsonian monkeys allows for the formation of a greater number of synapses and/or synapses of a greater area, which could help increasing the tonic GABAergic tone of pallidal terminals upon VApc and CM neurons in parkinsonism.

Although it has long been established that the firing rate of GPi neurons is increased in parkinsonian monkeys (Wichmann et al., 1999; Soares et al., 2004) and PD patients (Hutchison et al., 1994), information on the effects of abnormally increased BG GABAergic output on thalamic firing rates remains scarce. In normal monkeys, the firing rate of basal ganglia-receiving areas in the thalamus is reduced after electrical stimulation of the GPi (Anderson et al., 2003). Similarly, high-frequency stimulation of the GPi in MPTP-treated monkeys resulted in lower firing rates in the ventral motor thalamus (Kammermeier et al., 2016). Studies in MPTP-treated monkeys have found increased burst firing in the motor thalamus (Guehl et al., 2003; Pessiglione et al., 2005), and similarly high levels of bursting have been shown in corresponding thalamic areas in PD patients (Zirh et al., 1998; Magnin et al., 2000; Molnar et al., 2005). In addition, neurons in basal ganglia-receiving areas of the thalamus display decreased firing rate in PD patients compared to recordings from healthy controls (Molnar et al., 2005). Furthermore, studies of MPTP-treated monkeys suggest an increased metabolic activity in the ventral motor thalamus (Mitchell et al., 1989; Rolland et al., 2007), possibly reflecting increased activity of basal ganglia inputs. It is also known that the mRNA expression of the α1 subunit of GABA-A receptors is decreased in basal ganglia-receiving areas of the thalamus in rodent models of PD (Caruncho et al., 1997; Chadha et al., 2000), which may result from unusually increased synaptic release of GABA in the parkinsonian condition. Altogether these findings indicate that the basal ganglia-mediated GABAergic regulation of the thalamocortical system is disrupted, and that these changes likely contribute to the pathophysiology of the basal ganglia-thalamocortical loops in PD. Although, they do not demonstrate a causal relationship, our findings suggest that neuroplastic changes in the synaptic organization of the pallidothalamic terminals system may be critical in mediating these effects. Along those lines, a recent study showed that an increased number of GABAergic synapses from single GPe terminals on STN neurons is associated with an increased strength of pallidosubthalamic synapses (Fan et al., 2012). Knowing that GPe and GPi GABAergic terminals share common ultrastructural features (large size, large number of mitochondria, multisynaptic) (Kultas-Ilinsky and Ilinsky, 1990; Ilinsky et al., 1997; Kultas-Ilinsky et al., 1997), neuroplastic changes in the number of pallidothalamic synapses may also be associated with changes in the strength of the pallidothalamic connections in parkinsonism.

An important issue that remains to be investigated is the compensatory versus maladaptive nature of these utrastructural changes and their association with the development and severity of parkinsonian motor signs. An assessment of the anatomical and functional integrity of the pallidothalamic system during the course of nigrostriatal dopamine denervation in the chronic MPTP-treated monkey model of PD (Masilamoni and Smith, 2018) is warranted to further address these issues.

### Mitochondrial morphology alterations in the pallidothalamic terminals of parkinsonian monkey

Mitochondria are critically important for proper synaptic function, due to their central role in ATP production, Ca^2+^ regulation, and other major signaling mechanisms. Presynaptic functions including neuronal activity and synaptic strength rely directly on mitochondria-driven ATP synthesis (Verstreken et al., 2005; Gulyas et al., 2006; Ivannikov et al., 2013; Kann et al., 2014; Rangaraju et al., 2014; Smith et al., 2016). Our data show that mitochondrial volumes in GPi terminals in VApc are larger than in CM, although there is no difference in mitochondrial number between the two nuclei. This finding suggests that GPi terminals in the VApc might have higher energetic demands than CM in order to maintain normal VApc firing (Justs et al., 2022). Furthermore, our 3D EM data demonstrate that there is a significant correlation between increases in mitochondrial volume, terminal volume, and synapses surface areas in both the VApc and CM of MPTP-treated parkinsonian monkeys. These observations suggest that the ultrastructure and the composition of presynaptic mitochondria might be associated with synaptic performance in both control and parkinsonian monkeys. Numerous studies have, indeed, demonstrated that mitochondrial volume changes correlate with firing rate, mitochondrial Ca^2+^ uptake and synaptic vesicle number in different brain regions (Kageyama and Wong-Riley, 1982; Wong-Riley, 1989; Gulyas et al., 2006; Ivannikov et al., 2013; Cserep et al., 2018; Lewis et al., 2018; Rodriguez-Moreno et al., 2018; Thomas et al., 2019).

Mitochondria from PD patients have been described as swollen with rounded rather than rod-like profiles with discontinuous outer membranes, reduced cristae and occasionally contain crystal-like inclusions (Trimmer et al., 2000). Studies in MPTP-treated animal models have also reported enlarged mitochondria with disordered mitochondrial cristae and electron dense inclusions (Tanaka et al., 1988; Mizukawa et al., 1990; Levi et al., 1994). In the present study, we did not notice any mitochondrial inclusion or disordered mitochondrial cristae in pallidal terminals that innervate VApc and CM of parkinsonian monkeys, but significant differences in the mitochondrial morphology between control and MPTP-treated monkeys were found i.e. increased MCI and mitochondrial volume density in the VApc and CM, respectively, and significant difference in the mitotype correlation graph. Higher MCI in VApc indicates that the mitochondrial shape has a more complex and higher surface area relative to volume (Brown et al., 2022). This eventually leads to increased O_2_ consumption and oxidative phosphorylation. A similar phenotype of increased mitochondrial complexity with aging was also observed in aging mouse skeletal muscle (Leduc-Gaudet et al., 2015), postulated to reflect compensatory hyper fusion response to stress, previously reported in cellular systems (Shutt and McBride, 2013). In contrast, mitochondrial volume density increase in the GPi terminals of parkinsonian monkeys might reflect mitochondrial abundance to increase ATP production and meet energy demand necessary to maintain the tonic increase in pallidothalamic outflow (Bereiter-Hahn and Voth, 1994; Prakash et al., 2017).

## Conclusions

Altogether, results of this study demonstrate that GABAergic pallidothalamic terminals are endowed with a high level of structural plasticity that may contribute to the development and maintenance of the abnormal increase in pallidal outflow to the thalamus in the parkinsonian state. Furthermore, the evidence for ultrastructural differences between GPi terminals in VApc and CM suggests that morphologically distinct pallidothalamic terminals may underlie specific physiological properties of pallidal inputs to VApc or CM neurons in normal and diseased states. The present data lays the foundation for future electrophysiological studies that will examine transmitter dynamics and postsynaptic responses to eventually elucidate the functional consequences of ultrastructural changes in terminal volume and the number and size of synapses of pallidothalamic terminals in Parkinson’s disease pathophysiology.

## Supporting information

Sub_Figure_1

## Acknowledgements

This work was supported by the NIH grant P50 NA098965 and NIH/ORIP grant P51OD011132. Thanks to Susan Jenkins for technical assistance with tissue processing. Thanks are also due to Emily Benson of Renovo Neural Inc, and now Lerner Research Institute’s 3DEM Core (Cleveland Clinic) for sample preparation and generating SBF/SEM datasets.

---

Supplementary Figure 1. Photomicrographs of TH-immunostained coronal sections at the level of the pre-commissural striatum (A–B), post-commissural striatum (C–D), and midbrain dopaminergic cell groups (E–F) of a control (left column) and a MPTP-treated (right column) monkey. <u>Abbreviations</u>: CA: caudate nucleus; GPe: globus pallidus, external segment; GPi: globus pallidus, internal segment; LC: locus coeruleus; PU: putamen; SNCd: substantia nigra compacta, dorsal tier; SNCv: substantia nigra, ventral tier; Th: thalamus; VTA: ventral tegmental area. Scale bars: A–D = 5mm, E-F = 2mm.

