## Supplementary figures and images for "Structural Plasticity of GABAergic Pallidothalamic Terminals in MPTP-treated Parkinsonian Monkeys: A 3D Electron Microscopic Analysis"

### Sub_Figure_1.jpg

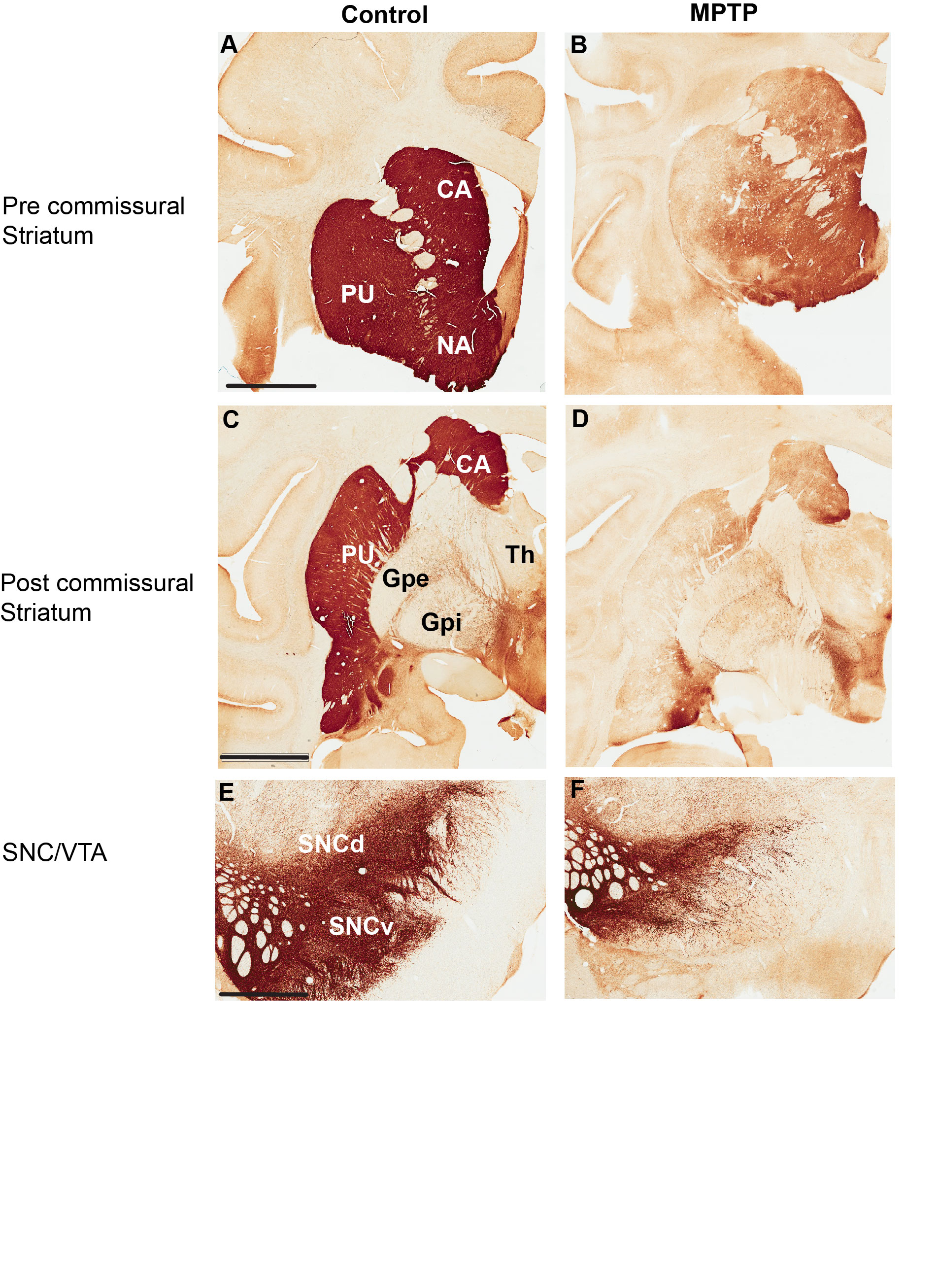
